# Machine Learning Prediction of Non-Coding Variant Impact in Cell-Class-Specific Human Retinal *Cis*-Regulatory Elements

**DOI:** 10.1101/2025.02.22.638679

**Authors:** Leah S. VandenBosch, Timothy J. Cherry

## Abstract

Non-coding variants in cis-regulatory elements such as promoters and enhancers contribute to inherited retinal diseases (IRDs), however, characterizing the functional impact of most regulatory variants remains challenging. To improve identification of variants of interest, we implemented machine learning using a gapped k-mer support vector machine approach trained on single nucleus ATAC-seq data from specific cell classes of the adult and developing human retina. We developed 18 distinct ML models to predict the impact of non-coding variants on 39,437 cell-class-specific regulatory elements. These models demonstrate accuracy over 90% and a high degree of cell class specificity. Variant Impact Prediction (VIP) scores highlight specific sequences within candidate CREs, including putative transcription factor (TF) binding motifs, that are predicted to alter CRE function if mutated. Correlations to massively parallel reporter assays support the predictive value of VIP scores to model single nucleotide variants and indels in a cell-class-specific manner. These analyses demonstrate the capacity for single nucleus epigenomic data to predict the impact of non-coding sequence variants and allow for rapid prioritization of patient variants for further functional analysis.

## Introduction

Inherited retinal disorders (IRDs) are a heterogeneous collection of diseases, affecting over 2 million individuals worldwide[1]. To date, over 300 genes have been implicated in IRDs (https://RetNet.org/)[2]. Many of these genes are expressed in just a subset of retinal cell types, shaping the genotype-phenotype correlation of each disorder [3, 4]. As many as half of all IRD cases cannot be explained by variants in protein-coding genes alone [5]. Well-studied disorders such as Blue Cone Monochromacy, Non-syndromic Congenital Retinal Non-Attachment, and Aniridia with Foveal Hypoplasia have demonstrated that non-coding genetic variants can cause retinal diseases[6-8]. It is likely that in additional cases without explicit genetic diagnoses, other non-coding variants could be contributing to disease. It is therefore important to characterize non-coding variants to improve upon our understanding of the genetic mechanisms underlying IRDs.

The vast majority of the human genome is composed of non-coding regions. This landscape contains *cis*-regulatory elements (CREs) such as promoters, enhancers, silencers, insulators and boundary elements, all of which shape the regulation of gene expression in cell-type-specific manners[9-12]. Genome-Wide Association Studies (GWAS) implicate non-coding genomic regions and CREs in retinal diseases such as Age-Related Macular Degeneration, Glaucoma, and Macular Degeneration [10, 13, 14]. Genetic variants within CREs have additionally been associated with altered retinal gene expression by expression quantitative trait locus (eQTL) analyses[15, 16]. Despite these analyses and associations, a lack of full non-coding characterization across complex tissues and limitations of existing tools contribute to a challenge in systematically interpreting individual variants within CREs, especially in the context of distinct cell classes.

The function of these CREs is determined through the complex interaction of DNA sequences and regulatory factor activity, in particular protein binding, such as that of transcription factors (TFs)[17-19]. These factors act together to regulate appropriate transcription and yield a complex epigenomic and expression profile for a given cell type[19]. DNA-protein interactions can be characterized by assays such as ATAC-seq— which identifies regions of genomic accessibility—and protein binding assays (ChIP-seq, CUT&RUN, CUT&Tag)[20, 21]. These assays can be used individually or together to identify and characterize candidate CREs. These identified elements are extremely diverse in accessibility, binding profiles, and activity, and may act in cell class and developmentally specific manners[22]. This makes the interpretation and characterization of such elements particularly challenging. Recent advancements have improved analyses by introducing single cell-resolution characterization[23]. However, in-depth characterization of non-coding elements remains a complex task.

Recently, machine learning based approaches have been trained with high-throughput - omics datasets to predict the inferred value or function of genetic sequences and variants, including on non-coding regulatory regions[24-26]. We have previously applied a gapped k-mer support vector machine (gkm-SVM/dSVM) based approach to retinal candidate CREs identified through bulk-tissue assays for DNA accessibility[27]. This approach was capable of distinguishing key features within retinal CREs in a tissue-specific manner.

However, as the retina is a heterogeneous tissue made up of numerous cell types [28], this model has limited applications in terms of disease modeling. In particular, in the case of less abundant cell populations in the retina, a bulk tissue-based approach would be unable to identify crucial sequences or differentiate between cell classes within the retina.

In this study, we applied a similar gkm-SVM/dSVM approach trained with our retinal single nucleus ATAC-seq (snATAC-seq) datasets[29] and an additional RPE ATAC dataset[30] to generate cell-class-aware models. We generated 18 models trained on high confidence peak calls. These models were determined to be 91-97% accurate and highly cell class specific. We additionally performed *in silico* saturation mutagenesis on holdout data to generate variant impact prediction (VIP) scores for 22,983 regions using deltaSVM. These VIP scores were able to distinguish the predicted impact in crucial sequences in a cell-class specific manner. To demonstrate functional relevance, we compared the predictions of our model against bulk and single cell Massively Parallel Reporter Assays (MPRAs), demonstrating a high degree of accuracy in model VIP scores predicting variant impact. Further, we generated a series of predictions on IRD-associated GWAS hits in cell classes of interest. This analysis will allow us to focus on relevant cell classes for IRDs and prioritize disease-associated variants in future analyses.

## Methods

Code used in for pre-processing data, training models, generating VIP scores, and comparing to MPRAs can be found at github.com/TCherryLab/snATAC-ML-2025/.

### Training data

Retinal snATAC-seq data generated by our lab is available through GEO (GSE183684) and described in Thomas et al 2022[29]. Macular RPE data was obtained from GEO (GSE99287) and is described in Wang et al 2018 [30]. Pseudobulk peak calls derived from individual cluster calls were performed as previously described[29]. Macular RPE ATAC-seq fastqs were used to call peaks as previously described[27]. Summits defined in by macs2 were extended +/-150 bp to generate putative CRE regions. Peaks were sorted by macs2 peak score, andretinal cell classes were compared to each other to determine universally accessible regions likely used for housekeeping gene regulation. Universally accessible regions were removed from the peak sets, and the top 25k summits selected from the remaining peaks were used for model generation. For selection of training data, regions for each cell class were shuffled, and 20k regions were randomly selected, with remaining peaks set aside as for later use as outgroup test sets. For a uniform test set, all 80% training sets were cross-compared against the outgroup sets, to ensure no model was trained on outgroup data. Negative training data was selected from non-accessible background genomic regions and GC-matched to the positive training data as previously described[27].

### Model Training

Models were trained with ls-gkm gkmtrain[31] as previously described. Validation and sequence scoring similarly follows as previously described[27].

### TF motif identification

Vocabulary sequences and GWAS variants were scored for sequence motifs via the JASPAR PFM database (10^th^ release)[32]. Motif lineplots were generated through the identification of genomic locations of specific motifs within the Homer database or using scanmotifgenomewide from Homer[33]. For further detail on motif analysis, see http://github.com/TCherryLab/snATAC-ML-2025/

### MPRA Analysis comparison

Bespoke dSVM scoring for individual models was performed to match specific SNV and indel variants tested by MPRAs in Zhao 2023[34]. For further detail on comparisons, see http://github.com/TCherryLab/snATAC-ML-2025/

## Results

### Models Built From snATAC are Accurate and Specific

Previous modeling from our group demonstrates the effectiveness of gapped k-mer Support Vector Machine (gkm-SVM) modeling on bulk ATAC-sequencing profiles of the human retina[27]. To create more granular predictions for unique cell classes of the retina, we utilized snATAC-seq data generated by our group[29]. These data were collected across three developmental timepoints and from adult human retinas, yielding 18 distinct clusters of developing and mature cell classes in the human retina (Figure 1A). Additionally, we integrated ATAC-seq data from adult macular retinal pigment epithelium (RPE) [30]. The top 25,000 ATAC peaks scored by macs2 from each of these clusters were used to train cell-class-specific gkm-SVM models. The peak sets were split such that 80% of the peaks were used for model training. The remaining 20% of peaks were cross referenced against the outgroups from other cell classes and combined to create a universal test set (Figure 1B).

**Figure 1.**
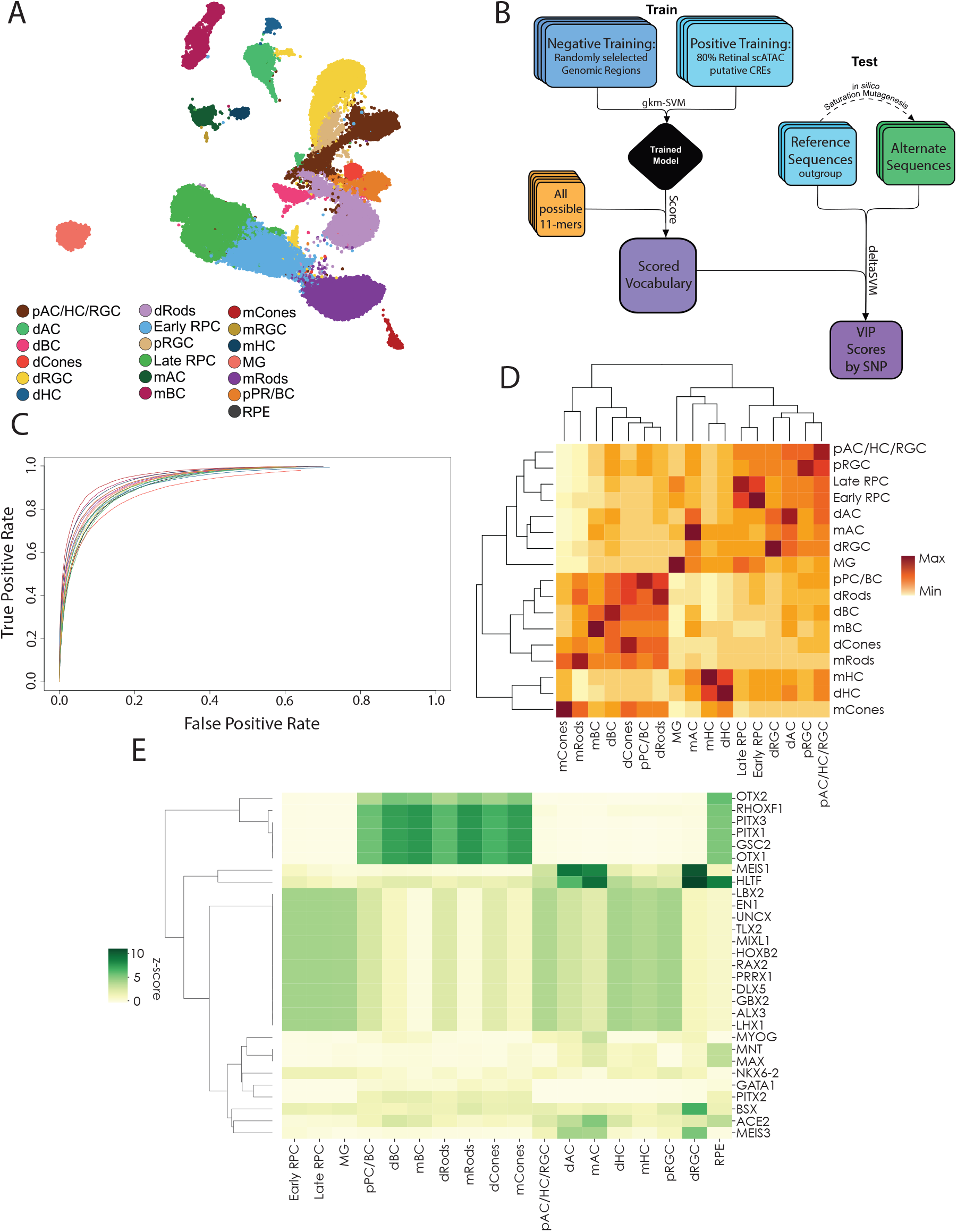
Generation of Cell Class-Specific ML Models A. UMAP of single nucleus(sn)ATAC-seqof developing and adult human retina, labeled by distinct cell classes[29] B. Model training paradigm for building gkm-SVM models from snATAC clusters in (A), and downstream analysis of variant sequences by deltaSVM. C. Receiver Operator Characteristic (ROC) curves from five-fold cross validation of all models built from cell classes as defined in (A). D. Model-to-model scoring of snATAC-defined CREs. Individual cell-class-specific models (columns) used to score CREs from individual cell class outgroup peaks (rows)with hierarchical clustering. E. Z-scores of top transcription factor family motifs (rows) present in the top 1% of scored 11mers by model (columns).

To demonstrate the accuracy of these models, the training data was used in five-fold cross validations of each model. These cross-validations were used to plot Receiver-Operator Characteristic (ROC) curves, mapping true and false positivity rates of each model (Figure 1C). All models performed extremely well, with Area Under the Curve (AUROC) values above 0.90 for each (Figure S1). To visualize model specificity, we scored holdout test peak sets for each cell class by each cell-class-specific model (Figure 1D). Comparing average scores for each peak set by model shows how models identify the most related sets of peaks, with self-recognizing-self as the highest scoring peak sets on average, as well as clustering of related cell classes. For instance, hierarchical clustering of cell class scoring by model gathers classes of rods, cones, and bipolar cells as well as precursors into related clusters, reflective of the similarity of accessible sequences and *cis*-regulation in each of these classes.

Using each model, a set of all possible 11-mers was scored for downstream applications. This scored set of vocabulary terms highlights the features driving each model. By subsetting the top 1% of scored vocabulary, we called known transcription factor (TF) binding motifs and identified which motifs are most highly enriched in the top vocabulary for each model. These motif calls were filtered by those TFs which are known to be expressed at any developmental stage of the human retina using snRNA-seq data[29] (Table S1). Using these motif counts, it is possible to highlight the top enriched motifs contributing to each model (Figure 1E).

### Variant Impact Predictions Identify Critical Sequence Motifs

To further testthe specificity of our newly generated models, we used them to predict the cell-type-specific consequences of disrupting consensus transcription factor binding motifs. First,using our 20% outgroup sequences, we performed *in silico* saturation mutagenesis and generated variant CRE sequences that contained all possible single nucleotide polymorphisms (SNPs). We then applied our models together with the deltaSVM pipeline [35] to generate Variant Impact Prediction (VIP) scores for each model for all possible mutations. Next, within the original hold out CREs, we identified all occurrences of specific consensus transcription factor binding motifs and averaged VIP scores for all possible variants within a +/-30bp window of each motif for each model. Scanning across the core sequence of a motif and its surrounding sequence context, we have demonstrated the cell-class-specific effect of motif disruption (Figure 2A-F). SNPs along the core motif of SOX2 and LHX2 demonstrate a high degree of specificity, with VIP scores reaching their most negative impact near the center of the motif, with a high degree of specificity for progenitor and Müller glial cells consistent with the expression of these TFs (Figure 2A&D). Similarly, TF motifs consistent with ONECUT family binding are scored most negatively in mature, developing and precursor horizontal cells (Figure 2B). TF motifs such as those compatible with OTX2 and CRX show even stronger VIP scores, with extreme specificity for bipolar and photoreceptor cell types and precursors the major cell classes that express these TFs (Figure 2C&F). In contrast, motifs of more general transcription factors expressed in many retinal cell types exhibit reduced specificity in model scoring predictions (Fig 2E). This is reflective of how each cell class utilizes these transcription factors, and thus functionally, disruption of a motif may be detrimental to one model but not the other. This trend is further demonstrated by the range of VIP scores by cell class along the core motif sequence for a wide variety of TF motifs (Figure S2A). While there are a large number of motifs for which disruption leads to strong VIP scores across multiple models (Figure 2G), many other motifs show highly specific, though potentially moderate effects (Figure 2H). VIP scores highlight still other motifs that are predicted to be more essential in mature cells than developing cells (Figure S2B). Altogether, these scores highlight the specificity and utility of these models to predict the cell-type-specific impact of regulatory variants..

**Figure 2.**
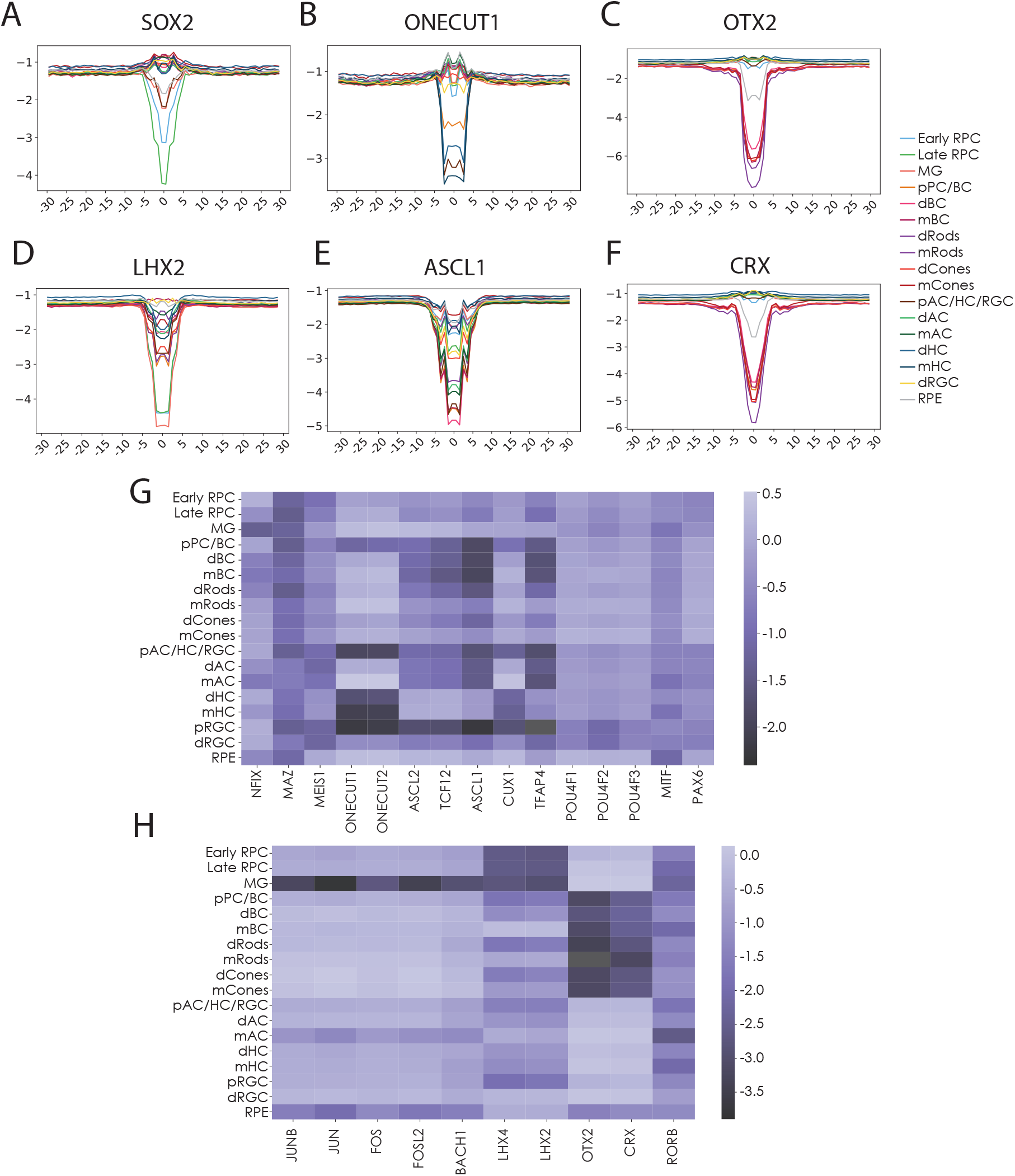
Cell-class-specific model scoring of transcription factor (TF) binding motif disruption. A-F. Line plots demonstrating average deltaSVM by cell class for *in silico* saturation mutagenesis of each base +/-25 bp around core TF motifs from outgroup of candidate CREs. G-H. Heatmaps of average Variant Impact Prediction (VIP) score for disruptions of motifs of interest (columns) by cell class (rows). G. High scoring contributors of transcription factor binding motifs. H. Moderate scoring contributors.

### Models Recreate CRE-Directed Expression Changes in Class-Specific Manner

We further assessed the predictive power of our models using existing data from functional studies of CRE variants. Zhao et al. characterized the impact of variants in CRX motifs within the mouse *Gnb3* promoter using a massively parallel reporter assay in single retinal cells. Characterization of these same variants with our models demonstrates that cell-class-specific VIP scores mirror changes in expression of reporter lines driven by this same promoter. Correlation of VIPs to variant reporter expression is highest in bipolar cells with a Pearson correlation of 0.575, with correlations of 0.49 in rod photoreceptors, while cell classes like Müller glial cells with low expression of *Gnb3* demonstrate little to no correlation between VIP scores and variant reporter expression (Figure 3B&C, Figure S3). This is consistent with higher expression of the gene in bipolar cells and rods, thus showing that the models are most predictive on elements active in the cell type of interest. These correlations between expression disruption and VIP are particularly clear in individual comparisons of individual and combinatorial deletions of CRX motifs from the *Gnb3* promoter sequence (Figure 3D&E). In each dataset, the second of five CRX motifs is identified as an extremely valuable sequence, with individual and combinatorial deletions of the sequence reflective of its importance to gene regulation. These correlations demonstrate the ability of these models to predict the impact of more complex variants beyond individual SNPs. Additionally, these results demonstrate the models’ capability for predicting the impact of variants in conserved loci across species, given that all models were trained on human CRE sequences, but *Gnb3* expression analysis was performed with the mouse promoter sequence. This is consistent with recent human-to-mouse comparisons of other tissues using gkm-SVM[36].

**Figure 3.**
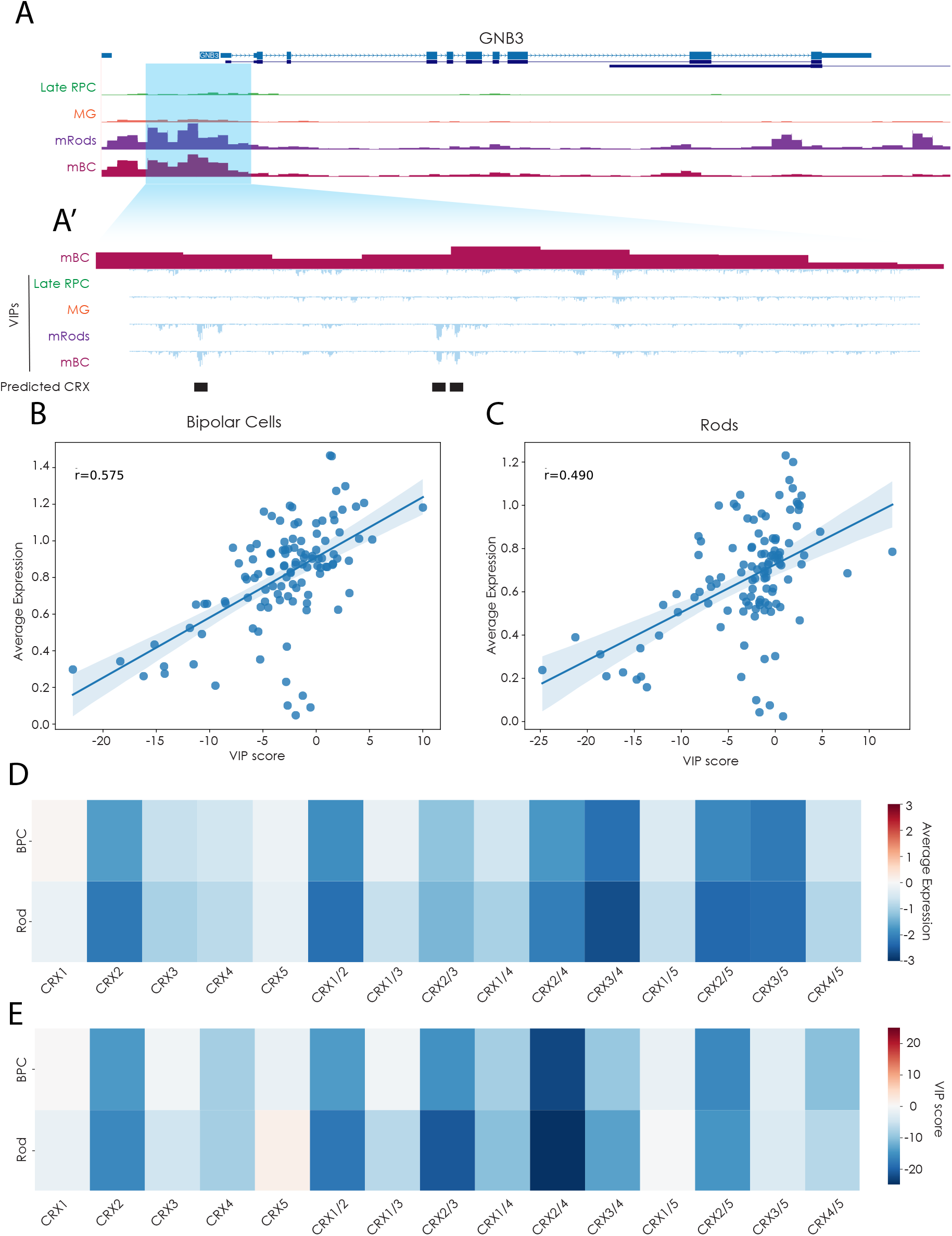
Cell-class-specific Variant Impact Prediction scores recapitulate disruptions modeled by massively parallel reporter assays. A. UCSC genome browser tracks of the *GNB3* locus and snATAC tracks for Late Progenitors, Muller Glia, Rods, and Bipolar cells [29]. A’ Zoomed view of highlighted promoter region with snATAC of Bipolar cells, and summed VIP scores for SNVs in region as scored by models for Late Progenitors, Muller Glia, Rods, and Bipolar cells. B,C. Scatter plots of correlation of Differential Expression and VIP scoring for disruptions of the *Gnb3* promoter. Average expression data from Zhao et al 2023. Expression and VIP scoring from B. Bipolar cells and C. Rods. r = Pearson correlation. D. Average expression values from massively parallel reporter assays of the *Gnb3* promoter with single and combinatorial CRX consensus binding motif deletions in Bipolar cells and Rod photoreceptors [34]. E. VIP scores for single and combinatorial CRX motif deletions as in D, scored using bipolar cell or rod photoreceptor-specific models.

### GWAS Predictions Narrow Variant Search Window

The value of these models becomes further apparent upon the intersection of Genome-Wide Association Studies and disease-associated variants. Studies of this nature have identified thousands of potential variants associated with complex retinal diseases, often concentrated in non-coding regions of the genome [14, 37]. Candidate regulatory variants can be narrowed down in scope based on many factors, such as DNA accessibility[14] (Figure 4A). However, this still leaves a daunting task of identifying which variants functionally contribute to phenotypes. By looking at sequence context and identifying candidate CREs where a disease-associated variant changes a predicted binding motif, the search for relevant variants can be further narrowed, but this still does not guarantee a functional change (Figure 4A).

**Figure 4.**
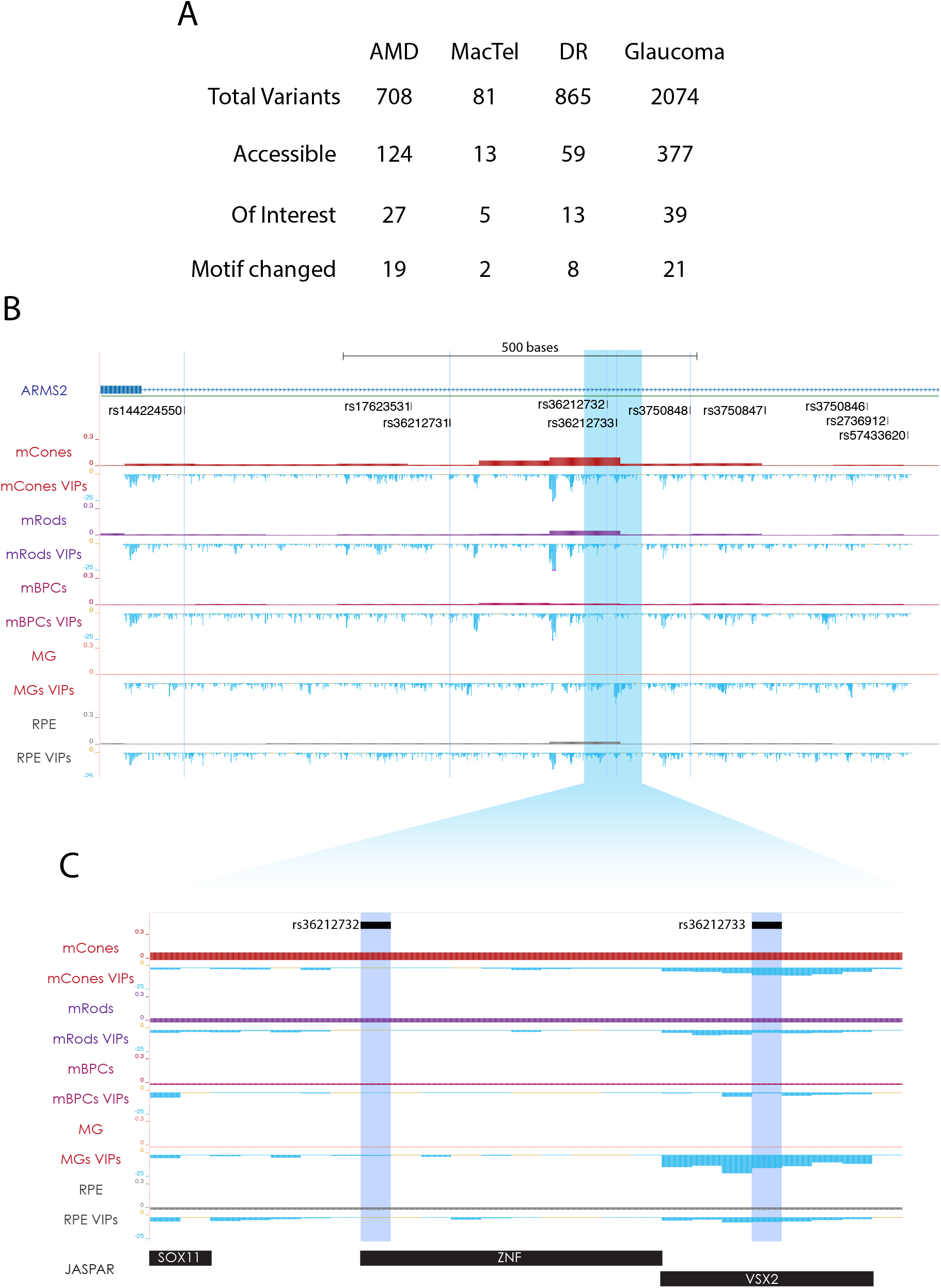
Prioritization of Variants of Interest from GWAS Studies. A. Variant filtration for GWAS identified variants [14]. Filtration by snATAC-derived accessibility in any cell class, accessibility in cell types of interest by retinal disorder, and presence in a change of motif at the site of variant. B. Tracks of snATAC-seq[29] and summed saturation mutagenesis VIP scores in region surrounding five variants identified for AMD (highlights). snATAC and VIP featured for Cones, Rods, Bipolar cells, Muller Glia, and RPE. C. Inset from (B). Specific scoring surrounding AMD-associated SNPs rs36212732 and rs36212733 and JASPAR-predicted transcription factor binding sites.

To predict which variants are most likely to have an impact on a cell type of interest, we generated VIP scores for all variants in an aggregated table of GWAS-identified sequence variants across four retinal diseases(Table S2). Using this scoring, we can identify regions of interest and variants with higher predicted impacts that are more likely to contribute to disease. Within the intronic region of ARMS2, there is a putative CRE which is accessible in photoreceptors, horizontal cells, and RPE. This region contains at least 5 variants associated with Age-Related Macular Degeneration (AMD) (Figure 4B). We have performed *in silico* saturation-level mutagenesis and generated VIP scores for all SNPs across the full accessible region. From this scoring, it is possible to identify regions of very negative VIP scores, indicating sequences where disruption may have a significant impact on CRE function. The scoring surrounding rs36212732 and rs36212733 in particular highlights the differences in VIP scores (Figure 4C). While both regions are accessible in cells of interest and fall within a predicted TF binding motif, only rs36212733 has a predicted impact. Both of these variants are highly associated with AMD, but VIP scoring indicates that only one likely has a strong impact on its own.

### A Resource for cell-class-specific Human Retinal Regulatory Variant Predictions

The analyses described above suggest that this cell-class-specific variant impact scoring strategy using the GKM-SVM/deltaSVM workflow trained on human retinal snATAC-seq data has several biologically relevant features. We therefore extended these scores to include a more inclusive set of 39,437 putative retinal CREs as defined by both our original ATAC-seq regions, as well as the top 10,000 peaks of retina-associated previously generated^38^ TF ChIP-seq data by MACS2 score. This analysis entitled “Cell-class-Specific Variant Impact Scores in Ocular Non-coding Sequences” (csVISIONS) is available on the UCSC genome browser to query and to compare with human retinal DNA-accessibility and gene expression(http://genome.ucsc.edu/s/CherryLab/csVISIONS_TrackHub). It is our hope that these predicted impact scores can assist other researchers in identifying and interpreting variants of interest within non-coding retinal regulatory elements.

## Discussion

The identification of causal non-coding variants in complex tissues presents a unique challenge in characterizing disease mechanisms and providing genetic diagnoses for IRDs. Variants have been identified across the genome in many non-coding regions, and the classification of a disease-associated variant as functional can be resource intensive, which is compounded when many variants must be tested across many cell types[10, 13]. In this study, we utilized single nucleus DNA accessibility data to train machine learning models on individual retinal cell classes. This level of resolution allows for distinct characterization of retinal cell classes and their regulatory mechanisms. We demonstrate that by using high-confidence DNA accessible regions, we are able to generate models that are both accurate, and cell-class-specific.

Comparisons of these models helps to demonstrates both their specificity and relatedness. As each model scores representative CREs and sequence variants, it is clear that each model values different sequences based on the sequence motifs that contribute to it. This can highlight the differences between each class, as well as relatedness between models that may have similar regulatory logic[38]. Hierarchical clustering of model scoring of peak sets demonstrates this cell class relatedness.

The driving features of each of these models can be discovered through explorations into the contributions of individual sequence motifs. In particular, TF binding motifs are highlighted by both scored model vocabulary, and in sequence disruption and VIP scoring. Through the summarization of TF motifs called in top scored 11mer vocabulary, we identify the motifs of TFs that contribute highly to each model. This can be observed by identifying calls for these sequences across the genome and comparing average VIPs for disruptions in and around those motifs. This identifies the TF motifs which generate strong-negative VIP scores when disrupted, and easily demonstrates the emergent features of models, as well as model relatedness based on combinations of motif VIP scoring.

It is necessary to ensure that these variant impact prediction scores are accurate. Comparisons to massively parallel reporter assays performed with cell-class-specific resolution [34] reveal a high degree of similarity between changes in reporter expression from sequence variants to VIP scoring of the same variants. This holds true in multiple cell types that express the construct. However, this scoring becomes less predictive when a construct is lowly expressed. This is to be expected, as major changes to expression are unlikely when the reference construct is lowly expressed if at all, as small variants are less likely to induce a high degree of expression. In cell classes where the reference construct is highly expressed, changes are more consistent and measurable. This is replicated in our models’ scoring of variants, which further identifies sequence motifs of interest that contributed most highly to the function of a CRE.

The value of this type of predictive scoring is apparent in larger screens for disease-associated variants. Many GWAS and eQTL studies have identified hundreds of variants associated with retinal diseases and expression of IRD-linked genes, many of which are present in non-coding sequences across the genome[14, 15, 39]. Prioritizing the risk of each variant becomes a challenge, as it is not always clear which sequence is likely to contribute to development or disease. Some filtration of these variants is possible through the identification of DNA sequence accessibility in cell classes of interest, and it is possible to identify if a TF motif is disrupted in the region. However, our models are able to quickly and efficiently screen variants of interest based on what degree they are predicted to disrupt a candidate CRE. By identifying variants of interest in this way, it is possible to streamline the characterization of variants in sequences of interest.

Ultimately, this study demonstrates that it is possible to create valuable, accurate models of *cis*-regulatory sequences in distinct cell classes of the retina. These models will be useful in future efforts to characterize cell class-specific regulation and identify non-coding variants of interest in disease modeling. Pre-screening of candidates will allow for researchers to narrow down candidate lists to reduce the complexity of future assays and focus on the most likely functional variants.

## Acknowledgements

This work was supported by grants from the BrightFocus Foundation M2022006F to L.S.V., the NIH National Eye Institute R01EY028584 and R01EY033364 and a gift from the Dawn’s Light foundation to T.J.C., and the Catalytic Collaboration Grant from the Brotman Baty Institute awarded to T.J.C. The authors thank Brendan McShane, LuLu Callies, Abbi Engel, and Theresa Chen for their review and valuable discussions regarding the progression of this manuscript.

## Author Contributions

L.S.V. and T.J.C. conceived of all experiments and analyses. L.S.V. developed and completed ML training and analysis pipeline. L.S.V. performed all final model training. L.S.V. performed all analyses. L.S.V. and T.J.C. wrote and revised the manuscript. L.S.V. and T.J.C. developed UCSC browser csVISIONS trackhub.

## Notes

http://genome.ucsc.edu/s/CherryLab/csVISIONS_TrackHub

http://github.com/TCherryLab/snATAC-ML-2025/

### Competing Interest Statement

The authors have declared no competing interest.

## References

1. Duncan, J.L., et al., Inherited Retinal Degenerations: Current Landscape and Knowledge Gaps. Transl Vis Sci Technol, 2018. 7(4): p. 6.

2. Network., D.S.R.R.I., A.a.h.s.u.e.r. Accessed, and January 18.

3. Schneider, N., et al., Inherited retinal diseases: Linking genes, disease-causing variants, and relevant therapeutic modalities. Prog Retin Eye Res, 2022. 89: p. 101029.

4. Lyu, P., et al., Gene regulatory networks controlling temporal patterning, neurogenesis, and cell-fate specification in mammalian retina. Cell Rep, 2021. 37(7): p. 109994.

5. Berger, W., B. Kloeckener-Gruissem, and J. Neidhardt, The molecular basis of human retinal and vitreoretinal diseases. Prog Retin Eye Res, 2010. 29(5): p. 335–75.

6. Ghiasvand, N.M., et al., Deletion of a remote enhancer near ATOH7 disrupts retinal neurogenesis, causing NCRNA disease. Nat Neurosci, 2011. 14(5): p. 578–86.

7. Nathans, J., et al., Molecular genetics of human blue cone monochromacy. Science, 1989. 245(4920): p. 831–8.

8. Bhatia, G., et al., Estimating and interpreting FST: the impact of rare variants. Genome Res, 2013. 23(9): p. 1514–21.

9. Howard, M.L. and E.H. Davidson, cis-Regulatory control circuits in development. Dev Biol, 2004. 271(1): p. 109–18.

10. Ward, L.D. and M. Kellis, Interpreting noncoding genetic variation in complex traits and human disease. Nat Biotechnol, 2012. 30(11): p. 1095–106.

11. Nord, A.S., et al., Rapid and pervasive changes in genome-wide enhancer usage during mammalian development. Cell, 2013. 155(7): p. 1521–31.

12. Perez-Cervantes, C., et al., Enhancer transcription identifies cis-regulatory elements for photoreceptor cell types. Development, 2020. 147(3).

13. Maurano, M.T., et al., Systematic localization of common disease-associated variation in regulatory DNA. Science, 2012. 337(6099): p. 1190–5.

14. Wang, S.K., et al., Single-cell multiome of the human retina and deep learning nominate causal variants in complex eye diseases. Cell Genom, 2022. 2(8).

15. Ratnapriya, R., et al., Retinal transcriptome and eQTL analyses identify genes associated with age-related macular degeneration. Nat Genet, 2019. 51(4): p. 606–610.

16. Strunz, T., et al., A mega-analysis of expression quantitative trait loci in retinal tissue. PLoS Genet, 2020. 16(9): p. e1008934.

17. Jindal, G.A. and E.K. Farley, Enhancer grammar in development, evolution, and disease: dependencies and interplay. Dev Cell, 2021. 56(5): p. 575–587.

18. Lee, T.I. and R.A. Young, Transcriptional regulation and its misregulation in disease. Cell, 2013. 152(6): p. 1237–51.

19. Spitz, F. and E.E. Furlong, Transcription factors: from enhancer binding to developmental control. Nat Rev Genet, 2012. 13(9): p. 613–26.

20. Klein, D.C. and S.J. Hainer, Genomic methods in profiling DNA accessibility and factor localization. Chromosome Res, 2020. 28(1): p. 69–85.

21. DeAngelis, J.T., W.J. Farrington, and T.O. Tollefsbol, An overview of epigenetic assays. Mol Biotechnol, 2008. 38(2): p. 179–83.

22. Buenrostro, J.D., et al., Single-cell chromatin accessibility reveals principles of regulatory variation. Nature, 2015. 523(7561): p. 486–90.

23. Preissl, S., K.J. Gaulton, and B. Ren, Characterizing cis-regulatory elements using single-cell epigenomics. Nat Rev Genet, 2023. 24(1): p. 21–43.

24. Holder, L.B., M.M. Haque, and M.K. Skinner, Machine learning for epigenetics and future medical applications. Epigenetics, 2017. 12(7): p. 505–514.

25. Chen, S., et al., DeepCAPE: A Deep Convolutional Neural Network for the Accurate Prediction of Enhancers. Genomics Proteomics Bioinformatics, 2021. 19(4): p. 565–577.

26. Zhou, J. and O.G. Troyanskaya, Predicting effects of noncoding variants with deep learning-based sequence model. Nat Methods, 2015. 12(10): p. 931–4.

27. VandenBosch, L.S., et al., Machine Learning Prediction of Non-Coding Variant Impact in Human Retinal cis-Regulatory Elements. Transl Vis Sci Technol, 2022. 11(4): p. 16.

28. Hoon, M., et al., Functional architecture of the retina: development and disease. Prog Retin Eye Res, 2014. 42: p. 44–84.

29. Thomas, E.D., et al., Cell-specific cis-regulatory elements and mechanisms of non-coding genetic disease in human retina and retinal organoids. Dev Cell, 2022. 57(6): p. 820–836 e6.

30. Wang, J., et al., ATAC-Seq analysis reveals a widespread decrease of chromatin accessibility in age-related macular degeneration. Nat Commun, 2018. 9(1): p. 1364.

31. Lee, D., LS-GKM: a new gkm-SVM for large-scale datasets. Bioinformatics, 2016. 32(14): p. 2196–8.

32. Rauluseviciute, I., et al., JASPAR 2024: 20th anniversary of the open-access database of transcription factor binding profiles. Nucleic Acids Res, 2024. 52(D1): p. D174–D182.

33. Heinz, S., et al., Simple combinations of lineage-determining transcription factors prime cis-regulatory elements required for macrophage and B cell identities. Mol Cell, 2010. 38(4): p. 576–89.

34. Zhao, S., et al., A single-cell massively parallel reporter assay detects cell-type-specific gene regulation. Nat Genet, 2023. 55(2): p. 346–354.

35. Lee, D., et al., A method to predict the impact of regulatory variants from DNA sequence. Nat Genet, 2015. 47(8): p. 955–61.

36. Oh, J.W. and M.A. Beer, Gapped-kmer sequence modeling robustly identifies regulatory vocabularies and distal enhancers conserved between evolutionarily distant mammals. Nat Commun, 2024. 15(1): p. 6464.

37. Montalbano, A., M.C. Canver, and N.E. Sanjana, High-Throughput Approaches to Pinpoint Function within the Noncoding Genome. Mol Cell, 2017. 68(1): p. 44–59.

38. Murphy, D.P., et al., Cis-regulatory basis of sister cell type divergence in the vertebrate retina. Elife, 2019. 8.

39. Fritsche, L.G., et al., A large genome-wide association study of age-related macular degeneration highlights contributions of rare and common variants. Nat Genet, 2016. 48(2): p. 134–43.

