## Supplemental Figure 1 for "Machine Learning Prediction of Non-Coding Variant Impact in Cell-Class-Specific Human Retinal *Cis*-Regulatory Elements"

A

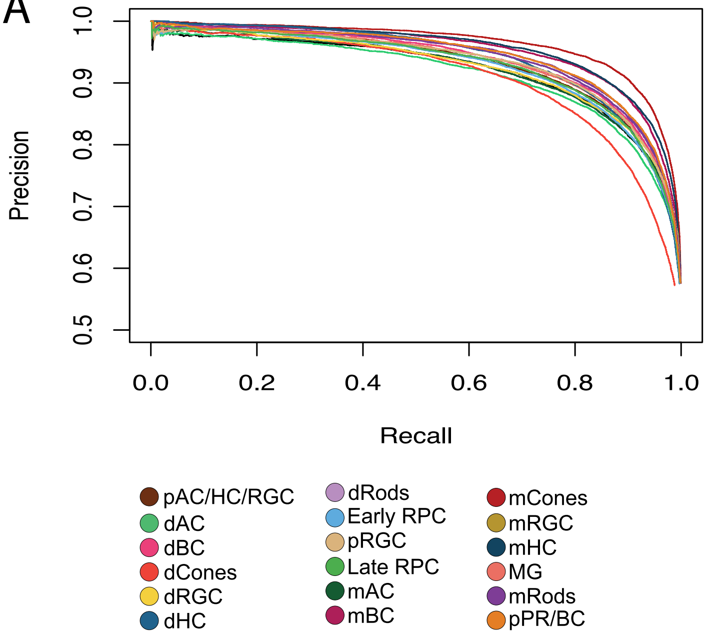


B

| Cell Class | ROC AUC | PRC AUC |
|---|---|---|
| Early RPC | 0.93 | 0.93 |
| pRGC | 0.94 | 0.94 |
| pAC/HC/RGC | 0.94 | 0.93 |
| dRGC | 0.93 | 0.92 |
| Late RPC | 0.93 | 0.93 |
| pPR/BC | 0.94 | 0.94 |
| dHC | 0.93 | 0.94 |
| dAC | 0.92 | 0.91 |
| dCones | 0.91 | 0.91 |
| dRods | 0.94 | 0.94 |
| dBC | 0.94 | 0.94 |
| pRGC | 0.94 | 0.93 |
| mHC | 0.95 | 0.95 |
| mAC | 0.93 | 0.92 |
| mCones | 0.96 | 0.96 |
| mRods | 0.94 | 0.94 |
| MG | 0.93 | 0.93 |
| mBC | 0.95 | 0.95 |
