## Supplementary figures and images for "Machine Learning Prediction of Non-Coding Variant Impact in Cell-Class-Specific Human Retinal *Cis*-Regulatory Elements"

### Supplemental Figure 2

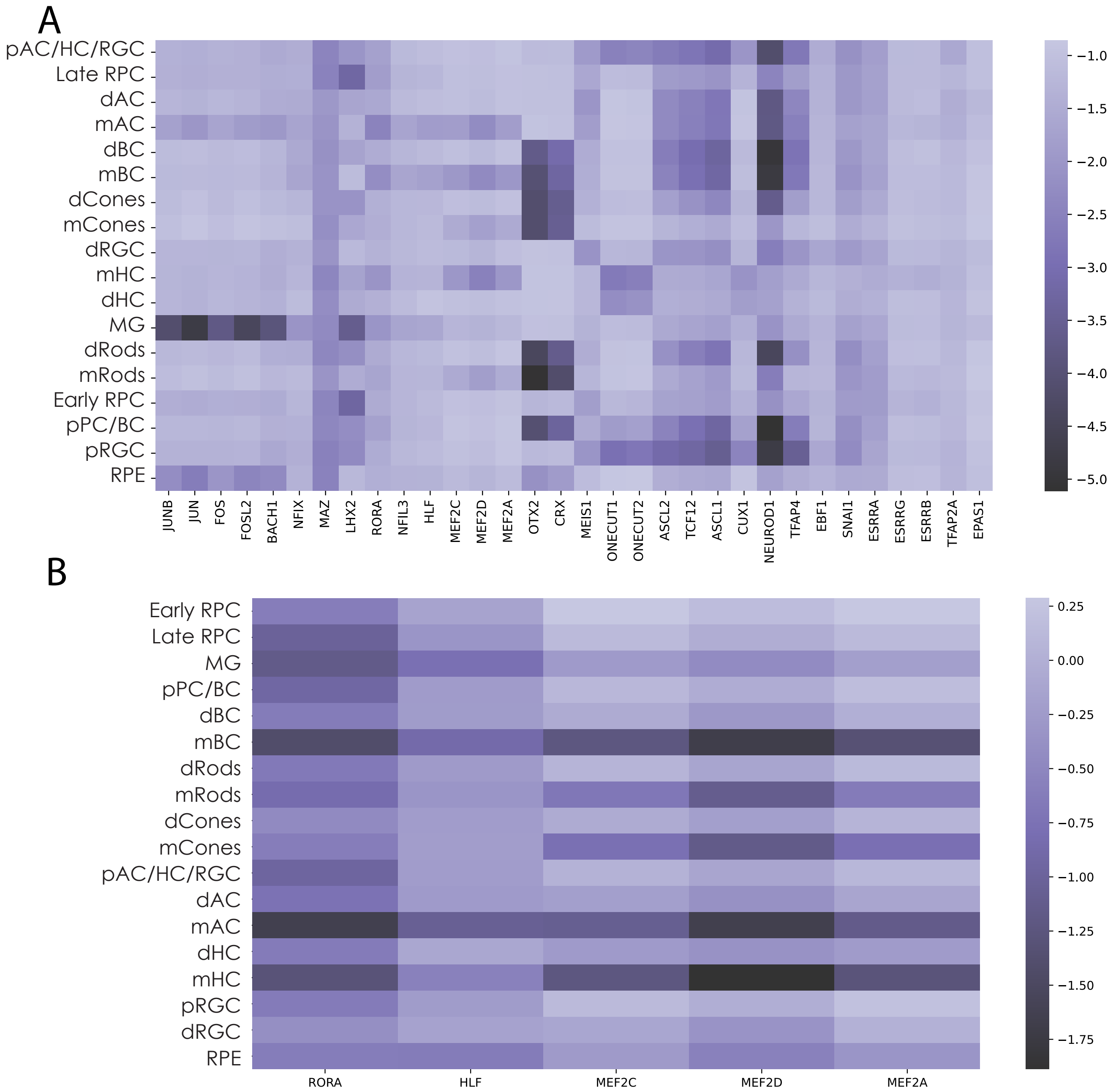

### Supplemental Figure 3

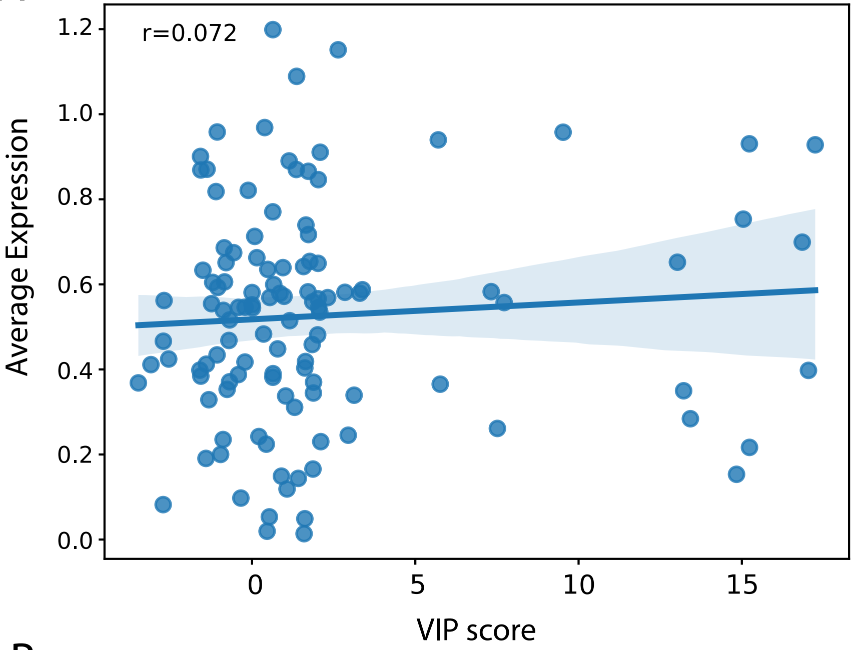

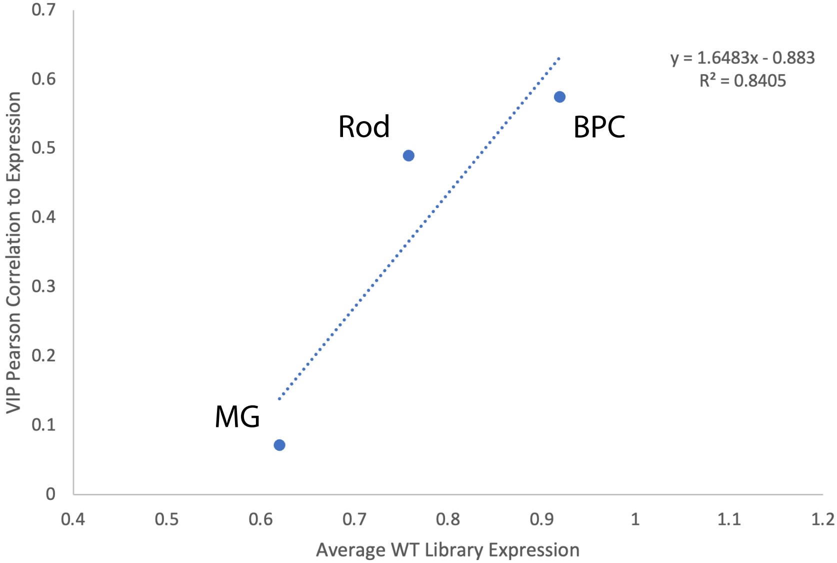
